# Large scale analysis of the diversity and complexity of the adult spinal cord neurotransmitter typology

**DOI:** 10.1101/518845

**Authors:** Andrea Pedroni, Konstantinos Ampatzis

## Abstract

The development of nervous system atlases is a fundamental pursuit in neuroscience, since they constitute a fundamental tool to improve our understanding of the nervous system and behavior. As such, neurotransmitter maps are valuable resources to decipher the nervous system organization and functionality. We present here the first comprehensive quantitative map of neurons found in the adult zebrafish spinal cord. Our study overlays detailed information regarding the anatomical positions, sizes, neurotransmitter phenotypes, and the projection patterns of the spinal neurons. We also show that neurotransmitter co-expression is much more extensive than previously assumed, suggesting that spinal networks are more complex than first recognized. As a first direct application of this atlas, we investigated the neurotransmitter diversity in the putative glutamatergic V2a interneuron assembly of the adult zebrafish spinal cord. These studies shed new light on the diverse and complex functions of this important interneuron class in the neuronal interplay governing the precise operation of the central pattern generators.

## INTRODUCTION

Neuronal networks in the spinal cord are able and sufficient to generate and control movements, and to receive and process sensory information (Arber, 2012; Goulding, 2009; Grillner and Jessell, 2009; Kiehn, 2016). Their functionality depends on the correct specification of different classes of neurons during development (Alaynick et al., 2011; Arber, 2012; Goulding, 2009; Jessell, 2000), which allows them to establish precise connections. Spinal neurons derive from specific progenitor pools in the spinal cord and express precisely a combinations of transcription factors (Alaynick et al., 2011; Arber, 2012; Goulding, 2009; Jessell, 2000). Their developmental diversification is well understood (Arber, 2012; Goulding, 2009; Jessell, 2000; Kiehn, 2016), but it is not clear how several functional characteristics of these cells are specified. A particularly important determinant of a neuron’s functionality is its neurotransmitter phenotype.

Neuronal communication involves the release and uptake of specific neurotransmitters (Rogawski and Barker, 1986; Schwartz, 2000) - endogenous chemical messengers used in intercellular signaling across synapses. The vertebrate nervous system uses neurotransmitters including glutamate, γ- aminobutyric acid (GABA), glycine, and acetylcholine to mediate biological functions such as sensory perception and to generate complex behaviors (Rogawski and Barker, 1986; Schwartz, 2000; Unwin, 1993). Neurons can be classified as excitatory, inhibitory, or modulatory based on their neurotransmitter phenotypes. Therefore, the adoption of a specific neurotransmitter system by a given neuron type defines its identity. To understand specific neurons’ roles in integrated neural networks, one must identify the transmitters they use to modulate their targets. Neuroanatomically precise maps of neurotransmitter typology distributions facilitate this by revealing correlations between the anatomical and functional neuronal architectures.

The zebrafish is an important model organism for high-throughput studies on neuronal circuits’ functions and behavior, and much is known about the different cell types in the zebrafish spinal cord (Ampatzis et al., 2013; Bernhardt et al., 1990, 1992; Björnfors and El Manira, 2016; Bradley et al., 2010; Böhm et al., 2016; Djenoune et al., 2017; Hale et al., 2001; Higashijima et al., 2004a; 2004b; Kimura et al., 2008; Liao and Fetcho, 2008; McLean et al., 2007; Menelaou et al., 2014; Satou et al., 2012; Stil and Drapeau, 2016). However, the number and identity of the spinal excitatory and inhibitory neurons that process sensory-related information are unknown, as are the neurotransmitter identities of the neurons that control and gate motor commands. This is a critical limitation because neuronal activity depends strongly on neurotransmitter identity. To overcome this limitation, we conducted the first systematic quantitative neurotransmitter phenotype analysis of neurons in adult zebrafish spinal networks by using an anatomical high-throughput strategy to investigate individual populations of spinal excitatory, inhibitory, and modulatory neurons. Our results reveal a previously unsuspected co-expression of different neurotransmitters in spinal cord neurons, and we show that these multi-phenotype neurons are far more numerous and widely-distributed in the spinal cord than previously assumed. We use this comprehensive neurotransmitter map to describe the co-existence of classical neurotransmitters in the presumed putative glutamatergic V2a interneuron population, revealing an unsuspected neurotransmitter co-expression within this cohered group of interneurons. The comprehensive neurotransmitter typology atlas presented here reveals an unforeseen diversity, complexity and dynamics in the principles which govern the structural organization of the adult zebrafish spinal cord and provides an anatomical framework to guide further functional dissection of spinal neuronal circuits.

## RESULTS

### Neuronal composition of the adult spinal cord

We first sought to determine the number of neurons in a representative hemisegment (segment 15) of the adult zebrafish spinal cord by using immunohistochemistry to detect the expression of the pan-neuronal marker HuC/D. This revealed that neurons were distributed throughout the adult spinal cord, from the most dorsal and medial part to the most lateral aspects (Figure 1A, 1B and 1D). However, only a small fraction of the labeled neurons was observed in the ventral part of the spinal hemisegment and the dorsal neuropil area (Figure 1D). Detailed quantification showed that an adult zebrafish spinal hemisegment contains 515.7 ± 8.865 neurons (segment 15; Figure 1C). While the soma sizes of the labeled spinal neurons varied considerably, the vast majority were small or medium-sized (41.17 ± 0.63 μm^2^, *n* = 2085 neurons; Figure 1E). These results show that the adult spinal cord has a well-defined and diverse neuron population, and provide a starting point for further characterizing the neurochemical architecture of adult zebrafish spinal cord networks.

**Figure 1.**
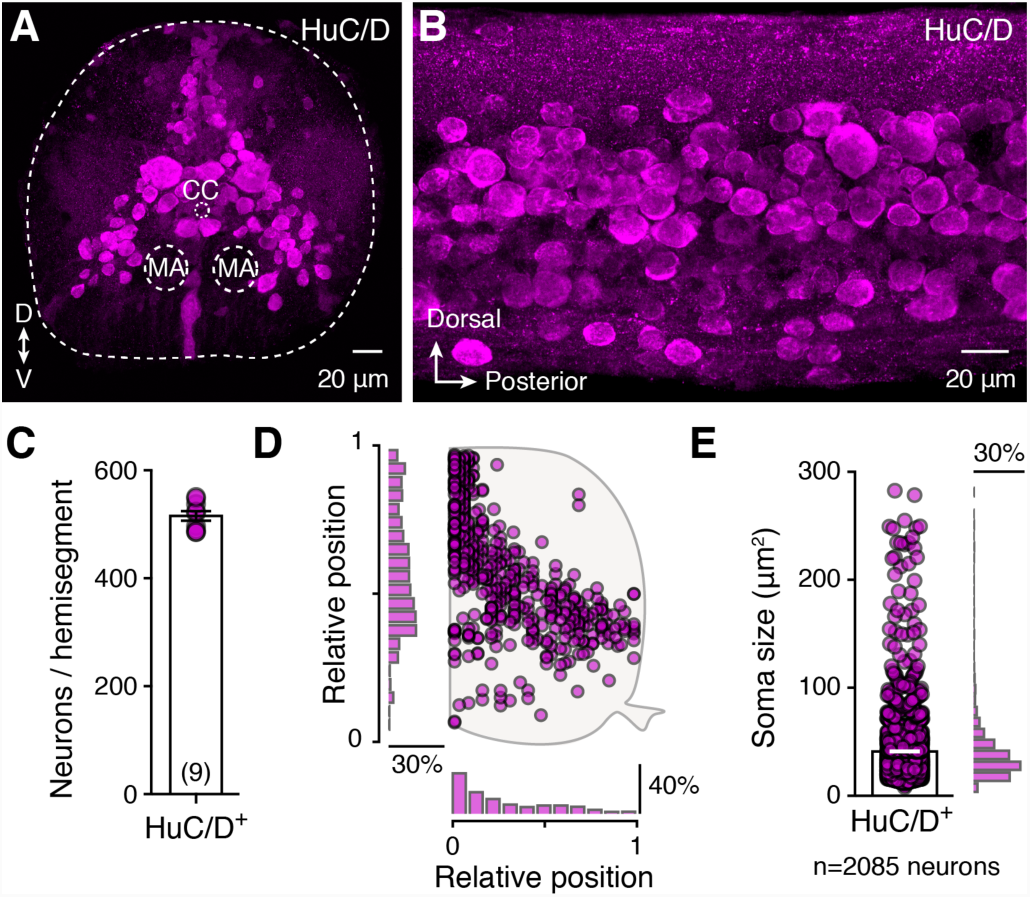
Neuroanatomy of adult zebrafish spinal cord. (A-B) Transverse section and whole mount adult zebrafish spinal cord showing the expression of the pan-neuronal marker HuC/D^+^neurons. (C) Quantification of the number of spinal neurons (HuC/D^+^) located in adult spinal cord hemisegment (segment 15). (D) Spatial distribution of the HuC/D^+^neurons with the medio-lateral and dorso-ventral density plot from one adult zebrafish spinal hemisegment (*n* = 478 labelled cells). (E) Quantification and distribution of the HuC/D^+^neurons soma size (*n* = 2085 neurons). Data are presented as mean ± SEM. CC, central canal; MA, Mauthner axon.

### Neurotransmitter typology of spinal cord neurons

Despite previous studies on zebrafish spinal neurotransmitter phenotypes (Higashijima et al., 2004a, 2004b), the number, size, and location of the neurons involved in the spinal networks are currently unknown. Therefore, to provide a reliable foundation for computational modeling and to identify new targets for electrophysiological recordings, we attempted to create a complete and detailed map of the neurotransmitter typology in the adult zebrafish spinal cord. All spinal neurons were found to express one of the classical neurotransmitters considered in this work (glutamate, GABA, glycine, acetylcholine and serotonin; Figure 2A-2D). In keeping with previous reports, we detected no dopaminergic or noradrenergic spinal neurons (McLean and Fetcho, 2004; Figure S1). The glutamatergic, GABAergic, and glycinergic neurons had similar distributions (Figure 2A-2C), while cholinergic neurons were almost absent from the dorsal part of the spinal cord (Figure 2D), and serotonergic neurons were only observed in the ventral part (Figure 2E). Quantification of individual neuronal classes revealed that most neurons are glutamatergic (212.1 ± 5.01 neurons, *n* = 9 zebrafish), GABAergic (145.5 ± 2.918 neurons, *n* = 10 zebrafish), and glycinergic (150 ± 3.179 neurons, *n* = 8 zebrafish; Figure 2F). Cholinergic neurons constitute a smaller population (79.78 ± 1.024 neurons, *n* = 9 zebrafish), and only few serotonergic neurons were found (11 ± 0.755 neurons, *n* = 7 zebrafish; Figure 2F). Finally, soma size measurements showed that all these neuronal populations had similar mean soma sizes, however the cholinergic and serotonergic neurons displayed the greatest and least soma size variability, respectively (glutamatergic: 35.11 ± 1.57 μm^2^, *n* = 206 neurons; GABAergic: 27.99 ± 0.525 μm^2^, *n* = 407 neurons; glycinergic: 41.04 ± 0.953 μm^2^, *n* = 379 neurons; cholinergic: 62.54 ± 3.427 μm^2^, *n* = 208 neurons; serotonergic: 29.5 ± 0.815 μm^2^, *n* = 37 neurons; Figure 2G and 2H). The distributions of the different neurotransmitter-expressing neurons in the adult zebrafish spinal cord are thus highly stereotypic and heterogeneous.

**Figure 2.**
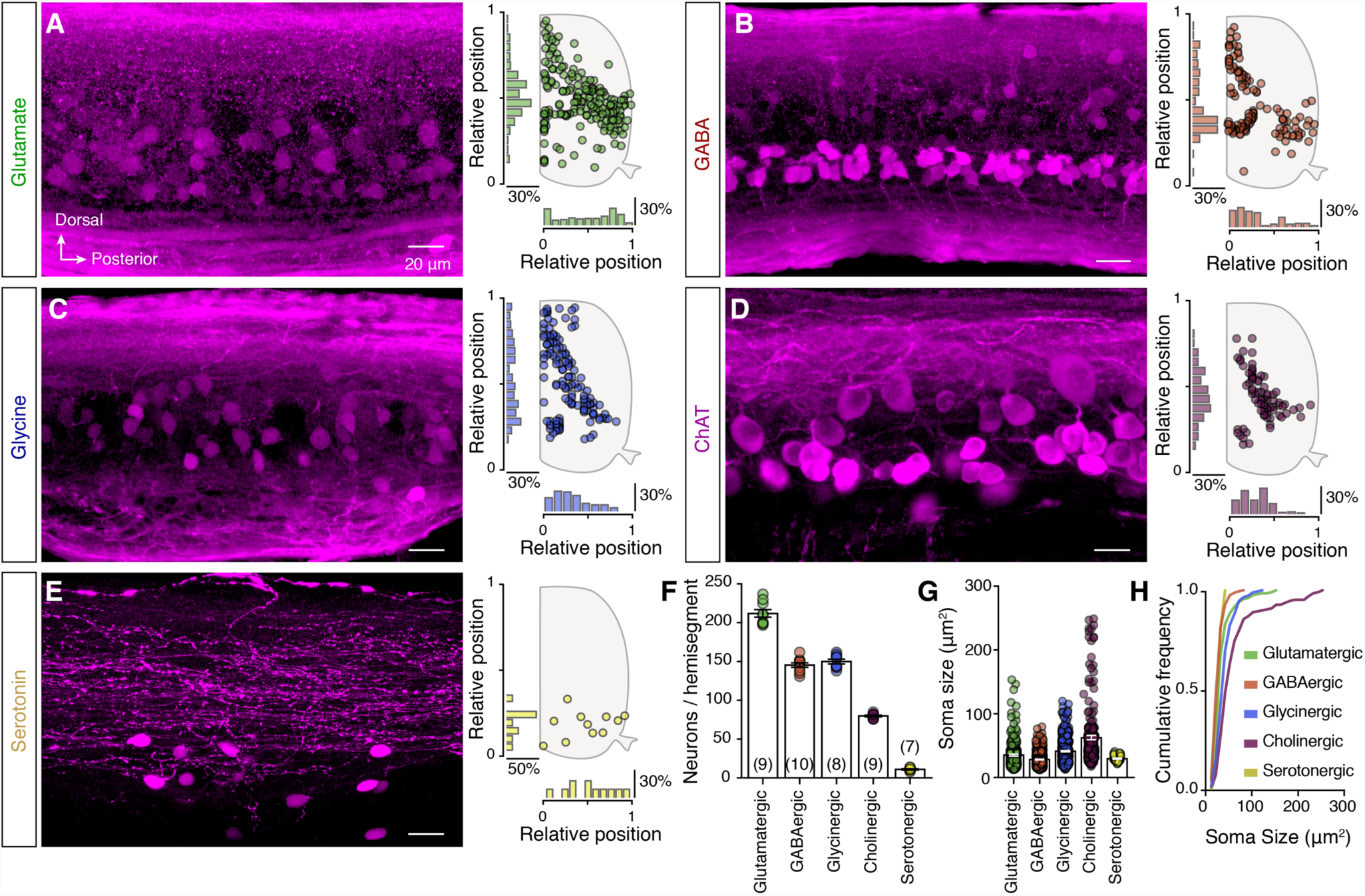
Neurotransmitter phenotypes of the adult zebrafish spinal neurons. (A-E) Representative whole mount photomicrographs showing part of the immunolabeled cells for glutamate, GABA, glycine, ChAT and serotonin, followed by a schematic representation of the spatial distribution with the corresponding medio-lateral and dorso-ventral density plots from a single adult zebrafish spinal cord hemisegment. (F) Quantification of the total number of the labeled neurons expressing a specific neurotransmitter phenotype. (G-H) Quantification and cumulative frequency of labeled neurons soma size. Data are presented as mean ± SEM. For related data, see also Figure S1 and S5.

### Neurotransmitter phenotypes of projecting spinal neurons

The projection patterns of the spinal cord neurons must be understood to explain their inputs to the circuits that process sensory information and control motor behaviors. We therefore used an anatomical tracing technique to determine the positions, number, and sizes of the projecting spinal neurons. Specifically, we identified every neuron located in hemisegment 15 that projects over 5 or more spinal segments to a rostral (ascending) or caudal (descending) spinal cord (Figure 3A and 3B). We found that most ascending neurons (∼75%) are located in the dorsal and medial part of the spinal cord, whereas the descending neurons are located in the motor column area (Figure 3C and 3D). Furthermore, the ascending neurons comprise a significantly smaller population (71.29 ± 2.212 neurons) than the descending neurons (87.38 ± 2.639 neurons; unpaired *t-*test: t = 4.594, df = 13, *P* = 0.0005; Figure 3E), and their soma sizes differ (ascending: 38.33 ± 1.106 μm^2^, *n* = 217 neurons; descending: 41.76 ± 0.894 μm^2^, *n* = 185 neurons; unpaired *t-*test: t = 2.354, df = 400, *P* = 0.0191; Figure 3F).

**Figure 3.**
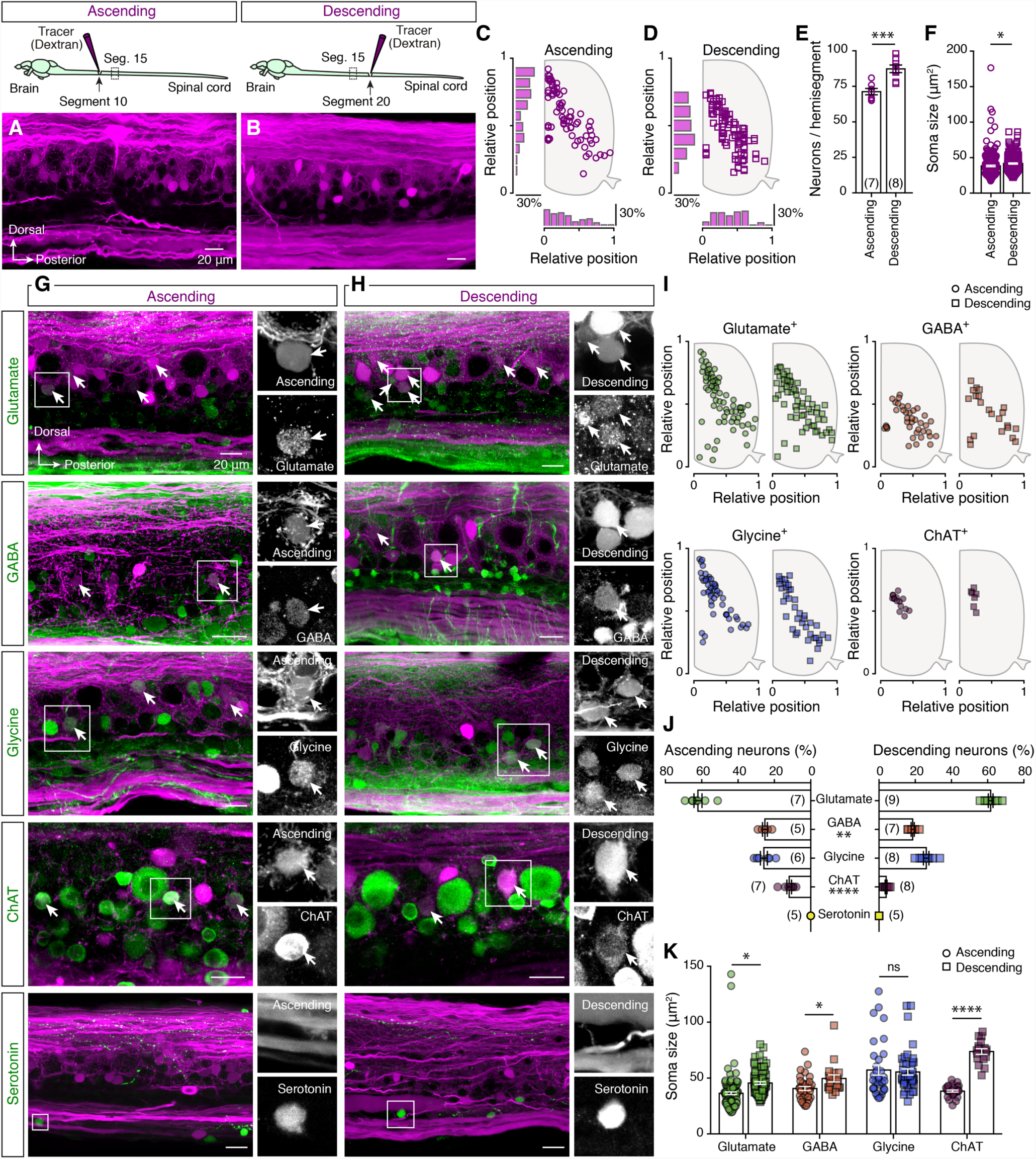
Neurotransmitter phenotype of projecting neurons. (A-B) Injection of a dextran tracer in segment 10 or 20 reveals the ascending and descending spinal projecting neurons, respectively, located in spinal cord segment 15. (C-D) Setting positions of the tracer-labeled ascending (circles) and descending (squares) neurons that project to the rostral or caudal part of the spinal cord revealed in one representative preparation. (E) Quantification of the total number of ascending and descending neurons detected in the spinal cord hemisegment. (F) Plot showing the soma sizes of the tracer-labeled ascending and descending neurons (Ascending: *n* = 217 neurons; Descending: *n* = 185 neurons). (G-H) Double staining between ascending or descending traced neurons (magenta) with glutamate, GABA, glycine, ChAT and serotonin (green). Arrows indicate the double labeled neurons. On the right sides there are single channel magnifications of the boxed area. (I) Spatial distribution of the ascending (circles) and descending (squares) traced neurons that express a specific neurotransmitter phenotype. (J) Quantification of percentage of tracer-positive ascending and descending projecting neurons expressing each neurotransmitter phenotype. (K) Soma sizes of the tracer-positive ascending (circles) and descending (squares) projecting neurons. Data are presented as mean ± SEM; \**P* < 0.05; \*\**P* < 0.01; \*\*\**P* < 0.001; \*\*\*\**P* < 0.0001; ns, non-significant.

To determine the projecting neurons’ neurotransmitter phenotypes, we combined the tracing with immunolabeling of the classical neurotransmitters (Figure 3G and 3H). This revealed that the ascending neurons located in the most dorsal area of the spinal cord are glutamatergic and glycinergic (Figure 3I), while GABAergic and cholinergic projecting neurons are co-distributed in the medial and ventral parts of the spinal cord (Figure 3I). In addition, no serotonergic neurons displayed ascending or descending projections extending over more than 5 segments (Figure 3G and 3H). Quantitative transmitter phenotype analyses showed that most projecting neurons are glutamatergic (ascending: 62.12 ± 2.2%, n = 7 zebrafish; descending: 61.67 ± 1.3%, *n* = 9 zebrafish; Figure 3J), while GABAergic (ascending: 25.29 ± 1.323 %, *n* = 5 zebrafish; descending: 18.6 ± 0.858 %, *n* = 7 zebrafish) and glycinergic (ascending: 25.87 ± 1.899%, *n* = 6 zebrafish; descending: 26.09 ± 1.542 %, *n* = 8 zebrafish) neurons form notably smaller populations. We also found a few projecting cholinergic neurons (ascending: 11.87 ± 1.297 %, *n* = 7 zebrafish; descending: 3.96 ± 0.506 %, *n* = 8 zebrafish; Figure 3J). With GABAergic (unpaired *t*-test: t = 4.45, df = 10, *P* = 0.0012; Figure 3J) and cholinergic neurons (unpaired *t*-test: t = 5.975, df = 13, *P* < 0.0001; Figure 3J) to exhibit a significant difference between the percentages of ascending and descending neurons. Our analysis suggests that similar patterns of excitation and inhibition are delivered to the rostral and caudal parts of the spinal cord. Finally, to determine whether different neuron types innervate the rostral and caudal parts of the spinal cord, we quantified the soma sizes of projecting neurons with respect to their neurotransmitter phenotypes (Figure 3K). Although in most cases the soma sizes of the ascending and descending neurons were significantly different (unpaired *t*-test: glutamatergic: t = 2.33, df = 143, *P* = 0.021; GABAergic: t = 2.652, df = 52, *P* = 0.01; Figure 3K), only the cholinergic neurons displayed populations with non-overlapping sizes (unpaired *t-* test: t = 15.22, df = 43, *P* < 0.0001; Figure 3K), suggesting that they constitute two distinct projecting subpopulations.

### Spinal neurons express multiple neurotransmitter phenotypes

Our analysis of neurotransmitter phenotypes in adult zebrafish spinal neurons suggested that the total number of neurons expressing a specific classical neurotransmitter is ∼600. Since we detected 515 neurons in each spinal cord hemisegment, this possibly implies that some spinal cord neurons (∼15 %) express multiple neurotransmitter phenotypes. To test this hypothesis, the extent of co-expression was determined using binary neurotransmitter immunodetection. We found several neurons co-express two neurotransmitter phenotypes (Figure 4A), and that these neurons have specific distribution patterns in the spinal cord (Figure 4C). Though, we found no co-expression of ChAT with serotonin and glycine, or of glycine with serotonin (Figure 4A). To determine whether neurons with dual neurotransmitter phenotypes comprise separate neuronal subpopulations that settle at distinct positions in the spinal cord, we measured the somas of double-labeled neurons (Figure 4B).

**Figure 4.**
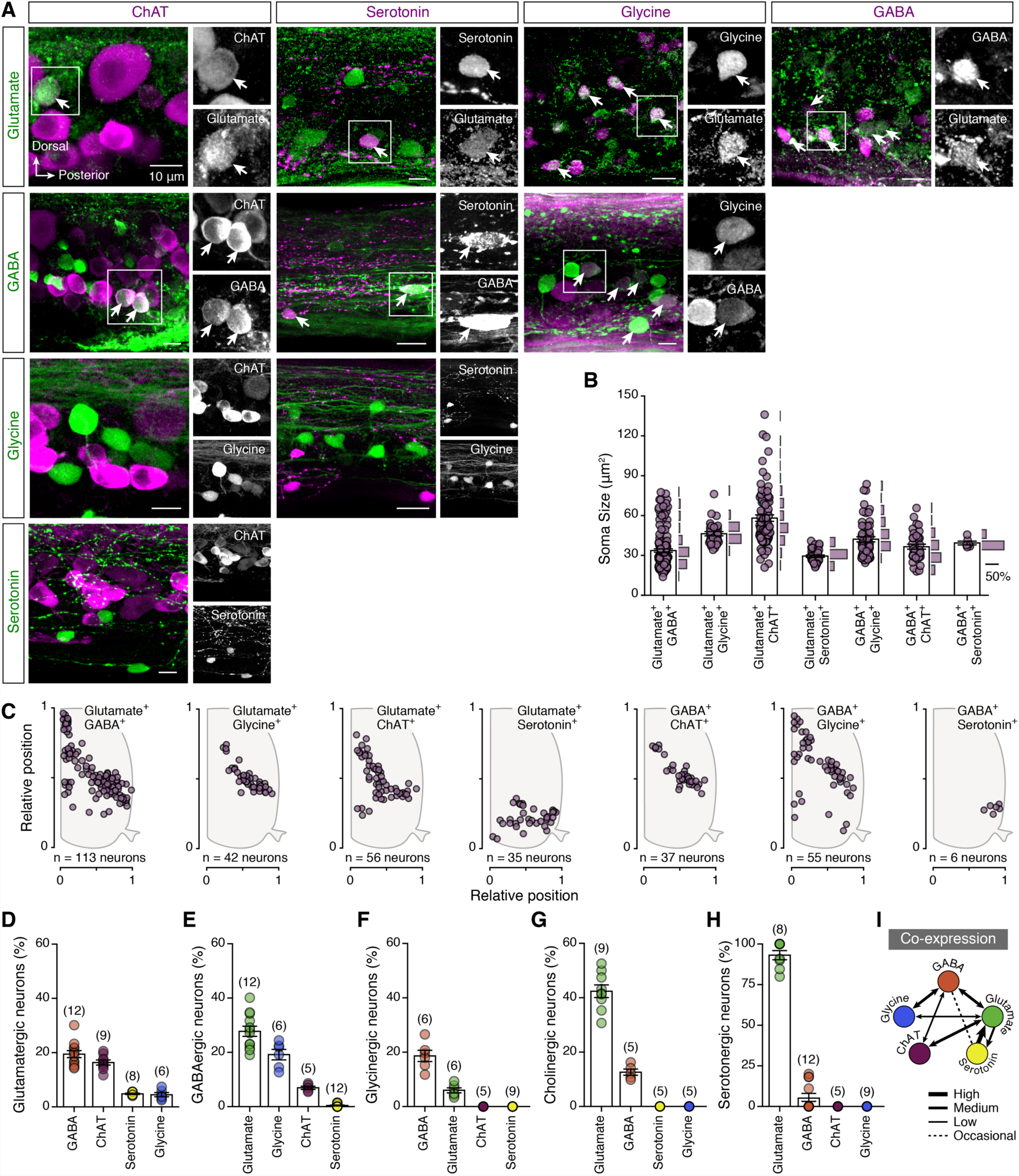
Spinal cord neurons express multiple neurotransmitter phenotypes. (A) Whole mount double immunolabeling between glutamate, GABA, glycine, ChAT and serotonin. In black and white are single channel images of the merged images. Arrows indicate the double labeled neurons. (B-C) Soma sizes and spatial distribution of the detected double-stained neurons in the adult zebrafish spinal cord hemisegment. (D-H) Quantification of percentage of glutamatergic, GABAergic, glycinergic, cholinergic (ChAT^+^) and serotonergic neurons that co-express other neurotransmitters. (I) Schematic relationship of the neurotransmitters co-expression from the adult zebrafish spinal cord neurons. Data are presented as mean ± SEM. For related data, see also Figure S2 and S3

Next, we determined the extent of dual neurotransmitter expression in different neuron populations. We found that a notable proportion of glutamatergic neurons are GABAergic (19.46 ± 1.302 %, *n* = 12 zebrafish), cholinergic (16.37 ± 0.927 %, *n* = 9 zebrafish), serotonergic (4.814 ± 0.224 %, *n* = 8 zebrafish), or glycinergic (4.507 ± 0.758 %, *n* = 6 zebrafish; Figure 4D). In addition, many GABAergic neurons co-express glutamate (27.76 ± 1.879 %, *n* = 12 zebrafish) or glycine (19.2 ± 1.922 %, *n* = 6 zebrafish), and a few were immunolabeled for choline acetyltransferase (ChAT; 6.965 ± 0.602 %, *n* = 5 zebrafish) or serotonin (0.413 ± 0.185 %, *n* = 12 zebrafish; Figure 4E). However, glycinergic neurons only observed to co-express GABA (18.67 ± 2.057 %, *n* = 6 zebrafish) and glutamate (6.008 ± 0.9372 %, *n* = 6 zebrafish; Figure 4F), as do cholinergic neurons (GABA: 12.62 ± 1.15 %, *n* = 5 zebrafish; Glutamate: 42.42 ± 2.311 %, *n* = 9 zebrafish; Figure 4G). Finally, many serotonergic neurons were found glutamatergic (93.19 ± 2.83 %, *n* = 8 zebrafish), and a few occasionally (4 out of 12 zebrafish) to co-express GABA (5.623 ± 2.47 %, *n* = 12 zebrafish; Figure 4H).

Most notably, the GABAergic/serotonergic neurons found to consist of a subpopulation of the serotonergic neurons that possess larger soma sizes (Figure 4B) and their distribution is restricted to the ventral and lateral part of the spinal cord (Figure 4C).

The validity of our observations was confirmed by immunohistochemistry and experiments using transgenic animal lines (*vGluT2a:GFP; Gad1b:GFP; GlyT2:GFP and Tph2:GFP*) to detect the proposed neurotransmitter phenotypes (*see* Experimental Procedures, Figure S2). To verify that spinal neurons can co-release different neurotransmitters, we performed *in situ* hybridization experiments using the sensitive RNAscope method to detect individual mRNAs for the vesicular glutamate transporter (vGluT2a, *slc17a6b*) found in neurons that release glutamate as a transmitter (Shigeri et al., 2004), the vesicular acetylcholine transporter (vAChT, *slc18a3b*; a specific transporter of cholinergic neurons, Weihe et al., 1996), and the vesicular GABA transporter (vGAT, *slc32a1*; also known as vIAAT – vesicular inhibitory amino acid transporter) a transporter for both GABAergic and glycinergic neurons (Chaudhry et al., 1998; Wojcik et al., 2006, Figure S3A – S3C). We observed the presence of different combinations (co-localizations) of the vesicular transporter mRNA puncta in individual neurons (Figure S3D – S3G), confirming that adult spinal cord neurons host the cellular machinery needed to store and release (co-transmit) different classical neurotransmitters. Interestingly, we also observed small populations of spinal neurons containing all three vesicular transporter mRNA puncta (Figure S3G), suggesting the existence of triple co-transmission. We verified this observation by immunohistochemistry, and investigated the distribution and soma sizes of spinal cholinergic neurons that co-express GABA and glutamate (Figure S3H).

Together, these data provide the first evidence that the characterization of neurons as being either excitatory or inhibitory is an oversimplification that does not properly reflect the neurotransmitter complexity of neuronal populations in the vertebrate spinal cord (Figure 4I).

### V2a interneuron neurotransmitter diversity: A proof-of-concept analysis

To evaluate our findings and the extent of neurotransmitter co-expression and dynamics, we performed a proof-of-concept analysis focusing on one of the most well-characterized spinal interneuron populations, the V2a interneurons (Arber, 2012; Goulding, 2009; Kiehn, 2016; 2011). V2a interneurons are one of the most important excitatory neuronal classes for the operation of the vertebrate locomotor network (Al-Mosawie et al., 2007; Crone et al., 2008; Dougherty and Kiehn, 2010; Hayashi et al., 2018; Joshi et al., 2009; Lundfald et al., 2007; Zhong et al., 2011), as demonstrated by studies on zebrafish (Ampatzis et al., 2014; Ausborn et al., 2012; Eklöf-Ljunggren et al., 2012; Kimura et al., 2006; McLean et al., 2008; McLean and Fetcho 2009; Menelaou et al., 2014; Song et al., 2018). In keeping with previous reports (Ampatzis et al., 2014), we detected 23.59 ± 0.503 V2a interneurons (*n* = 22 zebrafish; Figure S4B) per hemisegment in the adult zebrafish spinal cord. These interneurons were found to be distributed within the motor column (Figure S4C) and displayed variable soma sizes (Figure S4D). Detailed analysis of the neurotransmitter phenotype of the V2a interneuron population revealed that the vast majority (93.27 ± 1.116%, *n* = 17 zebrafish; Figure 5A and 5B) were glutamatergic. The glutamate^-^ V2a interneurons had significant smaller soma (24.61 ± 1.261 μm^2^) than those expressing glutamate (48.21 ± 2.902 μm^2^; unpaired *t-*test: t = 4.351, df = 104, *P* < 0.0001, Figure 5D). Interestingly, a smaller fraction of the V2a interneurons appeared to also express GABA (12.31 ± 1.217%, *n* = 9 zebrafish; Figure 5E and 5I), glycine (10.67 ± 0.84%, *n* = 7 zebrafish; Figure 5F and 5I), or ChAT (11.9 ± 0.755%, *n* = 13 zebrafish; Figure 5G and 5I). However, none were found to express serotonin (*n* = 6 zebrafish; Figure 5H and 5I). Moreover, the GABA^+^, glycine^+^and ChAT^+^V2a interneurons had distinct topographic distribution patterns (Figure 5J) and soma sizes (Figure 5K-5N), suggesting that they may constitute different subpopulations. Finally, we sought to determine whether the glutamatergic V2a interneurons could co-transmit these additional neurotransmitters by performing immunohistochemistry and *in situ* hybridization experiments to investigate their ability to produce the vesicular transporters for GABA and glycine (vGAT) and for acetylcholine (vAChT) (Figure S4). We detected vAChT, vGAT, and the glycinergic transporter (GlyT2) in presynaptic terminals (SV2^+^) of the V2a interneurons (GFP^+^, Figure S4E). In addition, vGAT or vAChT mRNAs were detected in a small proportion of the V2a interneurons by *in situ* hybridization (Figure S4F). These findings confirm our immunohistochemical observations (Figure 5E - 5N) and suggest that the V2a interneurons’ functional role in the organization and operation of the spinal cord networks controlling animals’ movements is more complex than previously recognized.

**Figure 5.**
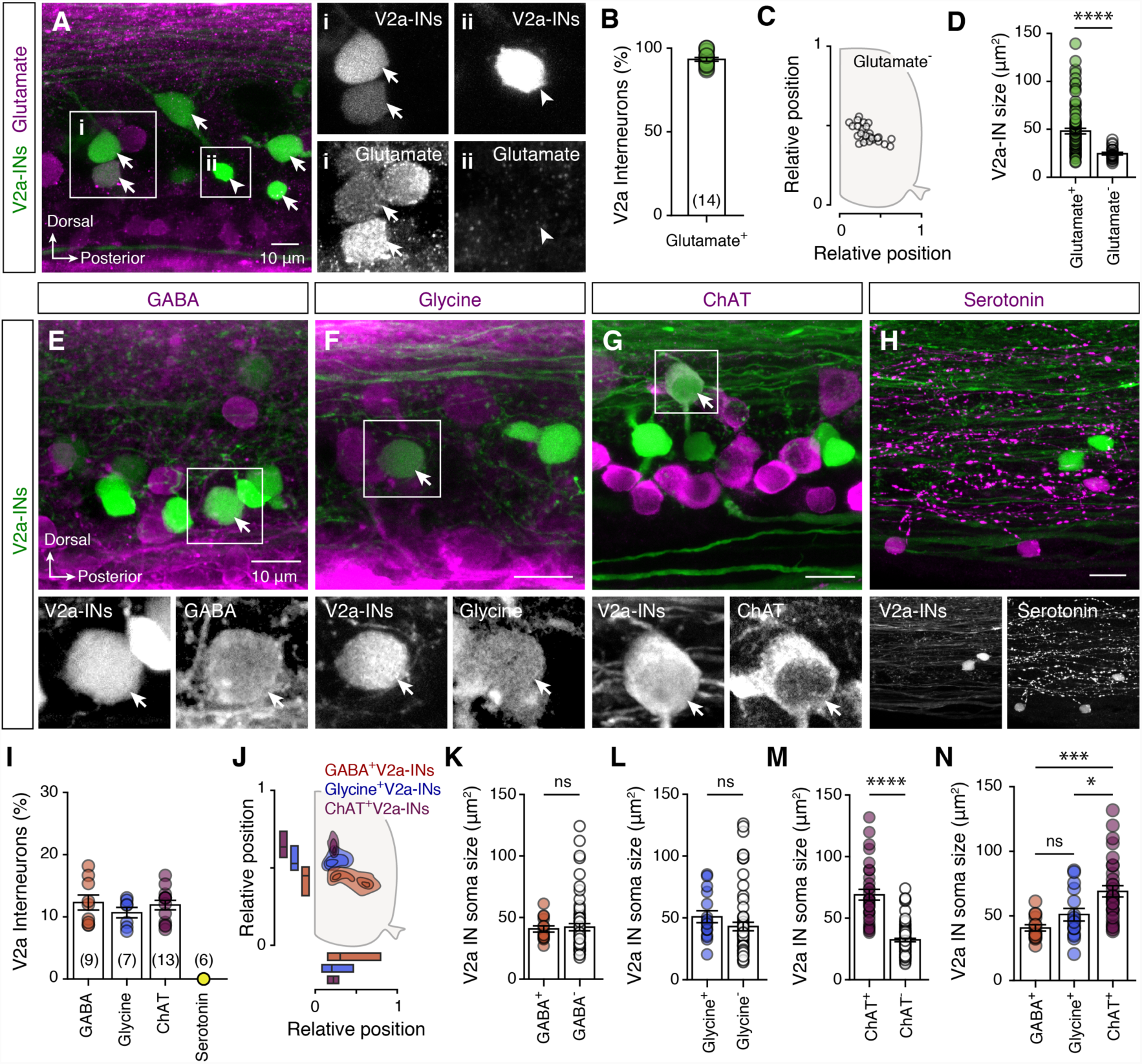
V2a interneuron neurotransmitter diversity. (A-B) Representative whole mount photomicrographs and analysis showing that the vast majority, but not all, of the adult zebrafish spinal cord V2a interneurons are expressing glutamate. Arrows indicate the double labeled neurons. Arrowheads indicate the non-glutamatergic V2a interneurons. (C) Setting positions of the glutamate^-^(open circles) V2a interneurons in the spinal cord. (D) Plot showing the difference in soma sizes of the glutamate^+^(green circles) and glutamate^-^(open circles) V2a interneurons. (E) Whole mount double immunolabeling between V2a interneurons with GABA, glycine, ChAT or serotonin. In black and white are single channel images of the merged images. Arrows indicate the double labeled neurons. (I-J) Analysis of the percentage and the topographic organization of the V2a interneurons that express GABA, glycine or ChAT. (K-M) Quantification of the V2a interneuron soma sizes that are immune-positive and immune-negative for the GABA, glycine or ChAT (unpaired *t-*test: t = 10.65, df = 111, *P* < 0.0001). (N) Comparison of the V2a interneuron soma sizes that express GABA, glycine or ChAT (one-way ANOVA: F(2,58) = 10.44, P = 0.0001). Data are presented as mean ± SEM. \**P* < 0.05; \*\*\**P* < 0.001; \*\*\*\**P* < 0.0001; ns, non-significant. For related data, see also Figure S4.

## DISCUSSION

We have conducted the first comprehensive classification of adult zebrafish neurons in a whole spinal cord hemisegment, revealing its total number of neurons, their sizes, the transmitter phenotypes they express, their setting positions, and their projection patterns. We have also established the extent of co-expression of the main classical neurotransmitters in spinal cord neurons, suggesting that the neurons’ chemical and anatomical organization is much more complex than previously recognized. Neuronal maps like that presented here, which describe distinct structural and biochemical features, provide essential guidance for future studies on the nervous system’s development and function. Cell-type specific neurotransmitter classifications of spinal neurons will enable further functional analyses of the diverse but stereotypic neuron populations that generate and gate sensory and motor functions to control animal movements.

Signal transmission in neuronal networks involves the release of neurotransmitters that bind specifically to membrane receptors on target neurons to mediate basic and complex biological functions. Since the identity of the neurotransmitters that a neuron synthesizes and releases is an important aspect of its differentiation fate, it is essential to understand the genetic programs that specify an individual neuron’s type and transmitter expression. The genetic programs that specify the spinal cord neuronal populations are well defined (Alaynick et al., 2011; Arber, 2012; Goulding, 2009; Jessell, 2000; Kiehn, 2016), but our understanding of neurotransmission within these neuronal classes is limited. Among the neurotransmitters of the nervous system, glutamate, GABA, glycine, acetylcholine, and serotonin are the most well-studied in the spinal cords of vertebrates (Alvarez et al., 2005; Antal et al., 1994; Brodin et al., 1990; Mahmood et al., 2009; Phelps et al., 1990; Pombal et al., 2001; Restrepo et al., 2009; Sueiro et al., 2004; Weber et al., 2007), including zebrafish (Barreiro-Iglesias et al., 2013; Bradley et al., 2010; Böhm et al., 2016; Higashijima et al., 2004a, 2004b; Liao and Fetcho, 2008; McLean and Fetcho, 2004). Several spinal interneuron types have been described in the developing zebrafish spinal cord based on their discrete morphological features (Bernhardt et al., 1990, 1992; Hale et al., 2001), which have been linked to specific neurotransmitter identities (Higashijima et al., 2004a, 2004b). These associations imply that most descending projecting interneurons express glutamate, while most ascending projecting neurons express GABA and/or glycine. This reinforces the notion that the principal descending input in the spinal cord is excitatory and the main ascending input is inhibitory. However, our tracing and immunodetection experiments suggest that similar numbers of inhibitory and excitatory neurons project to the rostral and caudal parts of the spinal cord, and the vast majority of these neurons are glutamatergic.

Our results firmly establish that many spinal cord neurons (∼15%; approximately 80-90 neurons) exhibit a multiple neurotransmitter phenotype. Views on neural function and synaptic communication have long been dominated by Dale’s principle (Eccles et al., 1954), which states that a neuron can only release a single neurotransmitter from all its synapses. This principle introduced a strongly reductionist approach to nervous system complexity by assigning each neuron to one of three functional classes (excitatory, inhibitory, or modulatory). Recently, however, several findings have complicated this simple characterization: there is growing evidence that neuronal populations in vertebrate and invertebrate nervous systems use multiple transmitter systems simultaneously. The possibility that neurons may release multiple neurotransmitters was first suggested by Burnstock (1976). Subsequent anatomical studies demonstrated the co-localization of multiple transmitters in single neurons (Hökfelt et al., 1977, 1987, 1998), and functional investigations have shown that many neuronal subtypes can store and release multiple neurotransmitters simultaneously (Granger et al., 2016; Hnasko and Edwards, 2012; Noh et al., 2010; Seal and Edwards, 2006; Vaaga et al., 2014). Nowadays, the concept of neurotransmitter co-release by single neurons is well accepted, and many, if not most, neurons are understood to use multiple transmission. However, the prevalence and physiological roles of co-transmission remain poorly understood, as is the synaptic circuitry involved.

The adult zebrafish spinal cord neurotransmitter atlas presented here is an essential resource for identifying currently unknown subpopulations of spinal neurons, and for future comparative studies on spinal circuit organization. Our anatomical mapping revealed a population of adult spinal neurons expressing both GABA and glycine, as previously demonstrated during zebrafish development (Higashijima et al., 2004a). It is well established that many neurons in the vertebrate spinal cord co-express and release these inhibitory neurotransmitters (Alvarez and Fyffe, 2007; Chery and De Koninck, 1999; Geiman et al., 2002; Taal and Holstege, 1994; Todd et al., 1996; Svensson et al., 2018). Moreover, in keeping with our observations here, it is well established that the vertebrate cholinergic spinal neurons (motoneurons) can coexpress and co-release glutamate along with acetylcholine (Bertuzzi et al., 2018; Meister et al., 1993; Mentis et al., 2005; Nishimaru et al., 2005). Interestingly, we also found that spinal cord neurons exhibit extensive co-expression of glutamate and GABA - two neurotransmitters with opposing functions. Although we did not investigate the release of these transmitters in this work, the possible co-release of glutamate and GABA from single nerve terminals in the brain has been demonstrated extensively (Beltrán and Gutiérrez, 2012; Galván and Gutiérrez, 2017; Noh et al., 2010; Root et al., 2014; Shabel et al., 2014; Yoo et al., 2016). Our findings support the existence of multi-transmitter neurons in the zebrafish spinal cord, as was already established in the lamprey spinal cord (Fernández-López et al., 2012) and the mammalian brain (Granger et al., 2017; Tritsch et al., 2016). However, the co-expression and co-release of these diverse transmitter combinations in mammalian spinal neurons has yet to be confirmed. Since the spinal cord is an evolutionarily conserved region of the central nervous system (Arber, 2012; Grillner, 2003; Grillner and Jessell, 2009), our results are probably relevant to organisms of higher phylogenetic order, including mammals. Based on this evolutionary perspective, we suggest that the diversity and complexity of zebrafish spinal neurons is likely to be echoed on larger scales in mammalian spinal systems, enabling better control of far more complex motor behaviors.

Our analysis also shows that the V2a interneurons form a functionally heterogeneous class of neurons that co-express GABA, glycine, or ChAT in addition to glutamate. Previous studies on the anatomical and functional organization of the V2a interneurons neglected the possibility that they might co-express and potentially co-release neurotransmitters other than glutamate (Ampatzis et al., 2014; Ausborn et al., 2012; Dougherty and Kiehn, 2010). However, earlier studies on zebrafish (Ampatzis et al., 2014; Ausborn et al., 2012; Song et al., 2018) and mice (Al-Mosawie et al., 2007; Zhong et al., 2011) clearly demonstrated that the V2a interneurons are functionally heterogeneous. In particular, the V2a interneurons in adult zebrafish form three discrete functional subpopulations that are incrementally recruited at different speeds of locomotion, and their recruitment pattern is not topographically organized (Ampatzis et al., 2014; Ausborn et al., 2012; Song et al., 2018). Although our findings indicate that a small fraction of the V2a interneurons can co-express other classical neurotransmitters in addition to glutamate, it seems very unlikely that these other neurotransmitters are released to control spinal motoneuron activity (Ampatzis et al., 2014; Song et al., 2016; Song et al., 2018). It seems more likely that that these additional neurotransmitters mediate the neuronal interplay needed for the precise operation of the central pattern generators, and may also contribute to the establishment of the necessary rostro-caudal delay.

Finally, neurotransmitter typology maps like that presented here are essential resources for clinical studies. Neurotransmitters are implicated in many neurological disorders, including schizophrenia, drug addiction, Parkinson’s disease, autism, Alzheimer’s disease, and attention deficit hyperactivity disorder (ADHD) (Abi-Dargham et al., 1997; Das et al., 2014; Huot et al., 2011; Márquez et al., 2017; Nakamura, 2013; Purkayastha et al., 2015; Rodríguez et al., 2012; Webster, 2001). Since neurotransmitter changes contribute to such diverse neurological disorders, comprehensive data on the diverse sets of neurons expressing specific neurotransmitters could enable the development of non-invasive therapies to restore nervous system function.

## Materials and Methods

### Experimental model

All animals were raised and kept in a core facility at the Karolinska Institute according to established procedures. Adult zebrafish (*Danio rerio*; *n* = 635; 9-10 weeks old; length: 16-19 mm; weight: 0.03-0.05 g) wild type (AB/Tübingen), and *Tg*(*vGluT2a:GFP; vGluT2a:DsRed; Gad1b:GFP; Gad1b:DsRed; GlyT2:GFP; Tph2:Gal4*^*y228*^*/UAS:GFP; Chx10:GFP*^*nns1*^) lines of either sex where used in this study. All experimental protocols were approved by the local Animal Research Ethical Committee, Stockholm (Ethical permit no. 9248-2017) and were performed in accordance with EU guidelines for the care and use of laboratory animals (86/609/CEE). All efforts were made to utilize only the minimum number of experimental animals necessary to produce reliable scientific data.

### Immunohistochemistry

All animals were deeply anesthetized with tricaine methane sulfonate (MS-222, Sigma-Aldrich, E10521). The spinal cords were then extracted and fixed in 4% paraformaldehyde (PFA) and 5% saturated picric acid (Sigma-Aldrich, P6744) in phosphate buffer saline (PBS) (0.01M; pH = 7.4, Santa Cruz Biotech., CAS30525-89-4) at 4°C for 2-14h. We performed immunolabeling in both whole mount spinal cords and in cryosections. For cryosections, the tissue was removed carefully and cryoprotected overnight in 30% (w/v) sucrose in PBS at 4°C, embedded in OCT Cryomount (Histolab, 45830), rapidly frozen in dry-ice-cooled isopentane (2-methylbutane; Sigma-Aldrich, 277258) at approximately – 35°C, and stored at –80°C until use. Transverse coronal plane cryosections (thickness: 25 μm) of the tissue were collected and processed for immunohistochemistry. In all cases (whole mount and cryosections) the tissue was washed three times for 5 min in PBS (0.01M; pH = 7.4, Santa Cruz Biotech., SC24946). Nonspecific protein binding sites were blocked with 4% normal donkey serum (NDS; Sigma-Aldrich, D9663) with 1% bovine serum albumin (BSA; Sigma-Aldrich, A2153) and 1% Triton X-100 (Sigma-Aldrich, T8787) in PBS for 1 h at room temperature (RT). Primary antibodies (Table S1) were diluted in 1 % of blocking solution and applied for 1-3 days at 4°C. After thorough buffer rinses the tissues were then incubated with the appropriate secondary antibodies (Table S1) diluted 1:500 or with streptavidin conjugated to Alexa Fluor 488 (1:500, ThermoFisher, S32354) Alexa Fluor 555 (1:500, ThermoFisher, S32355) or Alexa Fluor 647 (1:500, ThermoFisher, S32357) in 1 % Triton X-100 (Sigma-Aldrich, T8787) in PBS overnight at 4°C. Afterwards, the tissue was thoroughly rinsed in PBS and cover-slipped with fluorescent hard medium (VectorLabs; H-1400) on gelatine-coated microscope slides.

For the anti-GABA antisera raised in mouse (Table S1), the tissue was fixed in 4 % paraformaldehyde (PFA) containing 0.2 % glutaraldehyde (Sigma-Aldrich, G5882) and 5 % saturated picric acid in phosphate buffer saline (PBS; 0.01M; pH = 7.4) at 4°C for 10h. Finally, dual and triple neurotransmitter antibody stainings were performed in a sequential manner.

### Antibody specificity

The antisera in this study have been widely and successfully used previously in zebrafish to identify and describe the transmitter phenotype of neurons (anti-ChAT: Berg et al., 2018; Bertuzzi and Ampatzis 2018; Bertuzzi et al., 2018; Clemente et al., 2004; Mueller et al., 2004; Moly et al., 2014; Ohnmacht et al., 2016; Reimer et al., 2008; anti-GABA: Berg et al., 2018; Djenoune et al., 2017; Higashijima et al., 2004; Moly et al., 2014; Montgomery et al., 2016; Mueller et al., 2006; anti-Glycine: Berg et al., 2018; Moly et al., 2014; anti-Serotonin: Berg et al., 2018; Kuscha et al., 2012; McPherson et al., 2016; Montgomery et al., 2016; anti-TH: Ampatzis et al., 2008; Ampatzis and Dermon 2010; Rink and Wullimann, 2004; anti-DβH: Ampatzis et al., 2008), and the green fluorescent protein (anti-GFP: Barreiro-Iglesias et al., 2013; Böhm et al., 2016; Kuscha et al., 2012).

To evaluate the antibody specificity, adjacent sections or additional whole mount spinal cords were used in the absence of either primary or secondary antibody. In all cases no residual immunolabeling was detected. In addition, pre-incubation of the neurotransmitter antibodies used in this study with their corresponding antigen (100 μM - 400 μM; GABA (Sigma-Aldrich, A2129), glutamate (Sigma-Aldrich, G3291), glycine (Sigma-Aldrich, G6761), and serotonin (Sigma-Aldrich, 14927) for 1h at RT, eliminated any immunoreactivity.

To further confirm that our antibody immunodetections faithfully represented the neurotransmitter expression of spinal cord neurons, we performed a series of control experiments using transgenic animal lines (*vGluT2a, Gad1b, GlyT2 and Tph2*) combined with immunolabeling. We found that the vast majority or all of the neurons expressing the green fluorescent protein (GFP or DsRed) in either the vesicular glutamate transporter *vGluT2a*, a marker for glutamatergic excitatory neurons, the GABAergic inhibitory neuron marker *Gad1b*, the glycinergic neuronal marker *GlyT2* or the *Tph2* promoter for serotonergic neurons were immunolabeled with the anti-glutamate, anti-GABA, anti-glycine and anti-serotonin antibodies, respectively (Figure S5A-S5B). We also observed that immunostaining revealed more neurons than those expressing GFP (Glutamate^+^*vGluT2a*^*-*^: 26.77 ± 1.527 %, *n* = 8; GABA^+^*Gad1b*^*-*^: 13.59 ± 1.134 %, *n* = 8; Glycine^+^*GlyT2*^-^: 19.47 ± 0.449 %, *n* = 11; Serotonin^+^*Tph2*^-^: 23.66 ± 1.723 %, *n* = 7; Figure S5C), suggesting that although the transgenic zebrafish lines studied here consistently label the neurons of the expected neurotransmitter phenotype, they do not mark all neurons within a population. In addition, we used the transgenic animal lines as described above combined with single immunostaining for glutamate, GABA, glycine, ChAT and serotonin. We found that similar to dual immunodetections, we detected the vast majority of combinations of spinal co-expressing neurons (Figure S2). Regarding the specificity and reliability of the anti-glutamate immunodetection, we performed additional experiments to evaluate whether the glutamate^+^ cells are actually excitatory neurons, as glial cells can also store and release glutamate (Angulo et al., 2004; Hamilton and Attwell, 2010). Firstly, double staining with the anti-glutamate antisera and the NeuroTrace 500/525 Green fluorescent Nissl stain (ThermoFisher, N21480) showed that all the immunolabeled are indeed neurons (Figure S5D).

### RNAscope *in situ* hybridization

RNAscope *In situ* hybridization experiments shown in Figures S3 and S4, were performed according to manufacturer’s instructions in cryosections and in whole mound spinal cord preparations. The vesicular neurotransmitter transporter mRNAs were detected using RNAscope (Advanced Cell Diagnostics) zebrafish designed target probes (*slc17a6b*, also known as *vGluT2a*, Cat# 300031; *slc18a3b*, also known as *vAChT*, Cat# 300031-C2; *slc32a1*, also known as *vGAT* or *vIAAT*, Cat# 300031-C3). Following the *in situ* hybridization, cell nuclei and cell body were revealed using respectively the DAPI staining and anti-Elav 3+4 (HuC/D) immunodetection. In V2a interneuron (*Chx10:GFP*^*nns1*^), glutamate decarboxylase 1b transgenic (*GAD1b:GFP*), and in glycine transporter 2 (*GlyT2:GFP*) transgenic zebrafish line, the tissue was further processed for immunohistochemical detection of the GFP in the same way as in all immunodetections (see above, “Immunohistochemistry” section). The tissue was processed and mounted on Thermo Scientific™ Superfrost Plus™ Adhesion microscope slides and cover-slipped with anti-fade fluorescent mounting medium (Vectashield Hard Set, VectorLabs; H-1400).

### Ascending and descending neuron labeling

Zebrafish (*n* = 70) of either sex were anesthetized in 0.03% tricaine methane sulfonate (MS-222, Sigma-Aldrich, E10521). Retrograde labeling of ascending and descending spinal cord neurons located in the spinal segment 15 was performed using dye injections with biotinylated dextran (3000 MW; ThermoFisher, D7135) into segments 10 or 20 respectively. Afterwards, all animals were kept alive for at least 24h to allow retrograde transport of the tracer. Afterwards, all animals were deeply anesthetized with 0.1% MS-222. The spinal cords were dissected and fixed in 4% paraformaldehyde (PFA) and 5% saturated picric acid (Sigma-Aldrich, P6744) in phosphate buffer saline (PBS; 0.01M, pH = 7.4; Santa Cruz Biotech., CAS30525-89-4) at 4°C for 2-14h. The tissue was then washed extensively with PBS and incubated in streptavidin conjugated to Alexa Fluor 488 (dilution 1:500, ThermoFisher, S32354), Alexa Fluor 555 (1:500, ThermoFisher, S32355) or Alexa Fluor 647 (dilution 1:500, ThermoFisher, S32357) overnight at 4°C. Primary and secondary antibodies were applied as described before (see Experimental procedures, Immunohistochemistry section). After thorough buffer rinses the tissue was mounted on gelatine-coated microscope slides and cover-slipped with anti-fade fluorescent mounting medium (Vectashield Hard Set, VectorLabs; H-1400).

### Analysis

Imaging was carried out on a laser scanning confocal microscope (LSM 800, Zeiss) using the 40x objective. For the whole mount preparations, the whole hemisegment of the spinal cord (from lateral side to medial area of the central canal) was scanned generating a z-stack (z-step size = 0.5 – 1 μm). Cell counting was performed in hemisegment 15 of the adult zebrafish spinal cord (in whole mount preparations). The relative position of the somata of the neurons within the spinal cord, was calculated in whole mount preparations, using the lateral, dorsal, and ventral edges of the spinal cord as well as the central canal as landmarks. The relative position and soma sizes were measured using ImageJ (Schneider et al., 2012, NIH). For better visualization of our data, most of the whole mount images presented here were obtained by merging a subset of the original z-stack, showing the maximal intensity projections for each group. Analysis of the presynaptic terminals was performed of single plane confocal images taken using the 40x oil immerse objective. Colocalizing spots were detected by visual identification of structures whose color reflects the combined contribution of two or more antibodies in the merged image. In addition, analysis of fluorescent intensities for each channel was performed in Imagej software along single lines crossing the structures of interest (intensity spatial profile).

For the *In situ* hybridizations in cryosections (thickness: 25 μm) focal planes with homogeneous and reliable staining for each probe were taken generating a z-stack (z-stack 3-5 μm, z-step size < 1 μm). Only cells with visible DAPI^+^nuclei falling within the considered focal planes were examined for the presence of mRNA in the merged images. For the whole mount preparations, larger volumes (30 - 60 μm) of the spinal cord from lateral and from dorsal side ware scanned generating a z-stack (z-step size = 0.5 – 1 μm). As for the cryosections, only small subsets of the original stack (1-4 μm), with homogeneous and reliable staining and DAPI^+^nuclei falling within the considered focal planes, were examined for the presence of mRNA in the merged images. All the images were processed in Imagej software.

All figures and graphs were prepared with Adobe Photoshop and Adobe Illustrator (Adobe Systems Inc., San Jose, CA). Digital modifications of the images (brightness and contrast) were minimal so as not to affect the biological information. All double-labeled images were converted to magenta-green immunofluoresence to make this work more accessible to red-green color-blind readers.

### Statistics

The significance of differences between the means in experimental groups and conditions was analyzed using parametric tests two-tailed unpaired Student’s *t*-test or ordinary *one-way* ANOVA followed by *post hoc* Tukey multiple comparison test, using Prism (GraphPad Software Inc.). Significance levels indicated in all figures are as follows: \**P* < 0.05, \*\**P* < 0.01, \*\*\**P* < 0.001, \*\*\*\**P* < 0.0001. All data are presented as mean ± SEM (Standard error of mean). Finally, the *n* values reflect the final number of validated animals per group or the number of cells that were evaluated.

## ACKNOWELEDGMENTS

We thank Drs Shin-Ichi Higashijima, Emre Yaksi and Harold Burgess for sharing their transgenic animal lines. We also thank Drs. Mario Wullimann, Mark Masino, Konstantinos Meletis, Maria Bertuzzi and Luca Bertesaghi for their valuable discussion, comments, technical contributions to the project and assistance in preparing this manuscript. This work was supported by a grant from the Swedish Research Council (2015-03359 to K.A.), StratNeuro (to K.A.) Swedish Brain Foundation (FO2016-0007 to K.A.), Karolinska Institutet and Längmanska kulturfonden (BA17-0390 to K.A.).

## AUTHOR CONTRIBUTIONS

K.A. conceived the project and designed the experiments. A.P. and K.A. performed the experiments, analyzed the data, discussed the results, prepared the figures and wrote the manuscript.

