## Supplementary Figures for "Large scale analysis of the diversity and complexity of the adult spinal cord neurotransmitter typology"

**Andrea Pedroni and Konstantinos Ampatzis**
Karolinska Institutet, Department of Neuroscience, 171 77 Stockholm, Sweden

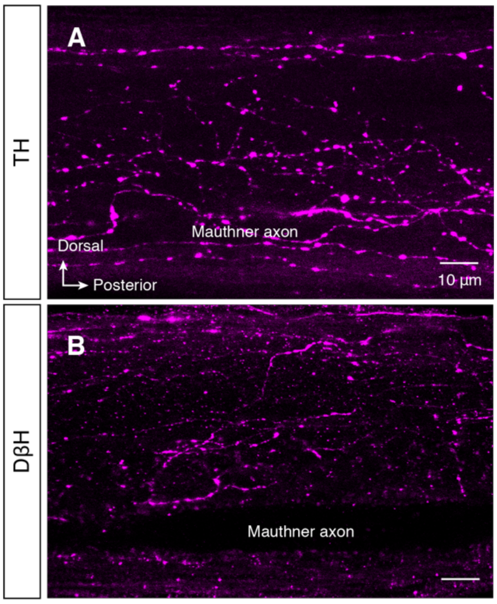


**Figure S1. Lack of spinal dopaminergic and noradrenergic neurons**
(A-B) Representative whole mount confocal images showing that only dopaminergic ($TH^+$) and noradrenergic ($D\beta H^+$) neuronal processes are detectable in the adult zebrafish spinal cord.

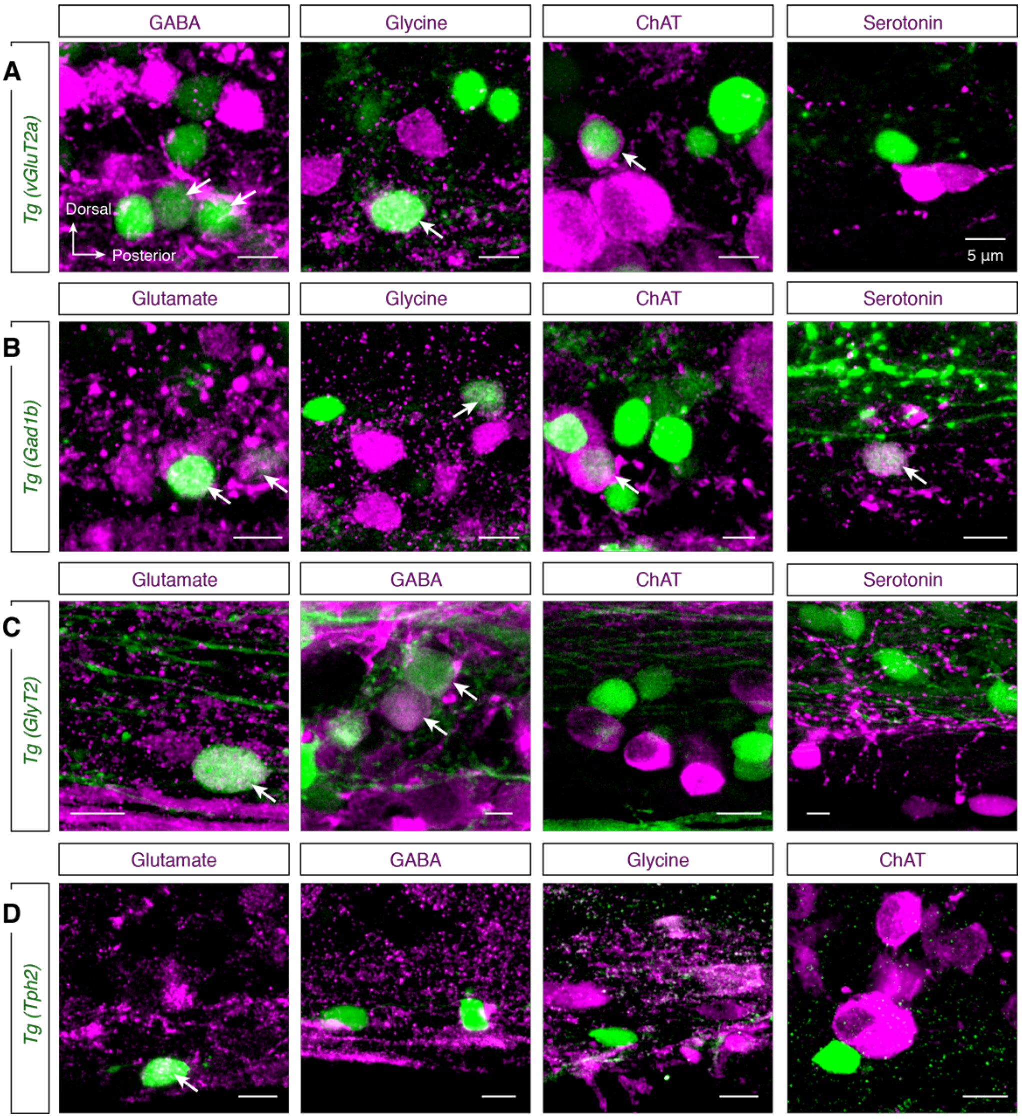


**Figure S2. The presence of co-expressing neurons in transgenic animal lines**
(A-D) Representative whole mount images from transgenic (*GlyT2*, *vGluT2a*, *Gad1b*, *Tph2*; green) adult zebrafish spinal cord preparations, immunolabeled for glutamate, GABA, glycine, ChAT and serotonin (magenta). Arrows indicate the double labeled cells.

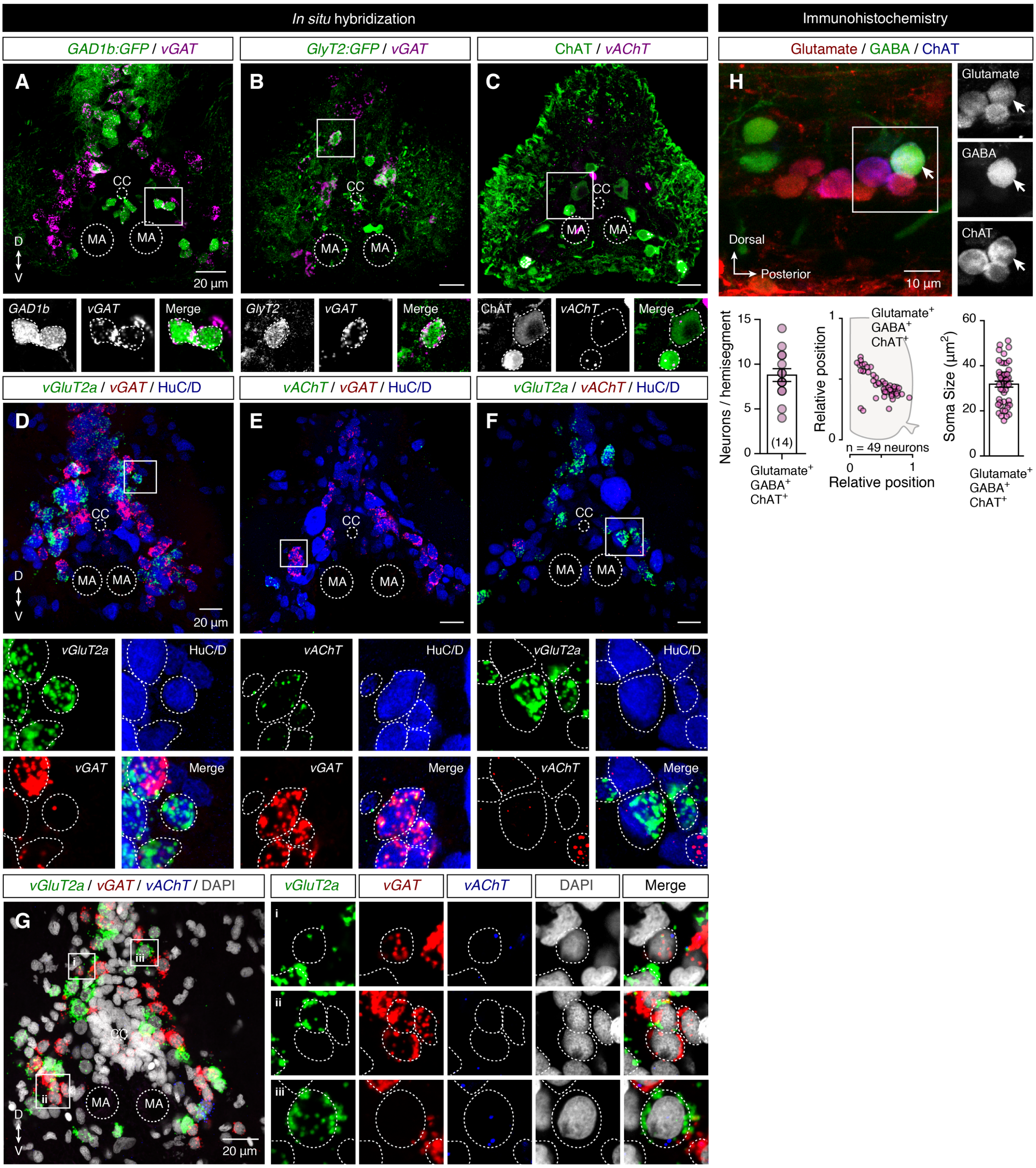


**Figure S3.** ***In situ*** **detection of the neurotransmitter's vesicular transporters**

(A-B) Confocal photomicrographs of transversal sections of the adult zebrafish spinal cord showing the presence of the *vGAT* mRNA in all the putative GABAergic (*GAD1b*) and Glycinergic (*GlyT2*) neurons.

(C) *In situ* hybridization reveals a small number of the vesicular acetylcholine transporter (*vAChT*) mRNA in all the cholinergic spinal neurons (ChAT$^+$).

(D-F) Co-localization of different vesicular transporter mRNAs in adult zebrafish spinal cord.

(G) Representative *in situ* hybridization for all the vesicular transporters (*vGluT2a*, *vGAT* and *vAChT*) mRNA showing a co-localization in the adult zebrafish spinal cord neurons.

(H) Triple immunolabeling confirms the existence of multi-expressing, glutamate (red), GABA (green) and ChAT (blue) neurons. Arrow indicates the triple labeled neuron. Quantification of the number and spatial distribution of cholinergic neurons co-expressing simultaneously glutamate and GABA in the adult zebrafish spinal cord hemisegment (segment 15). Quantification of the ChAT$^+$Glutamate$^+$GABA$^+$ neurons soma size.

Dotted lines represent the borders of the neurons. Data are presented as mean ± SEM. CC, central canal; MA, Mauthner axon.

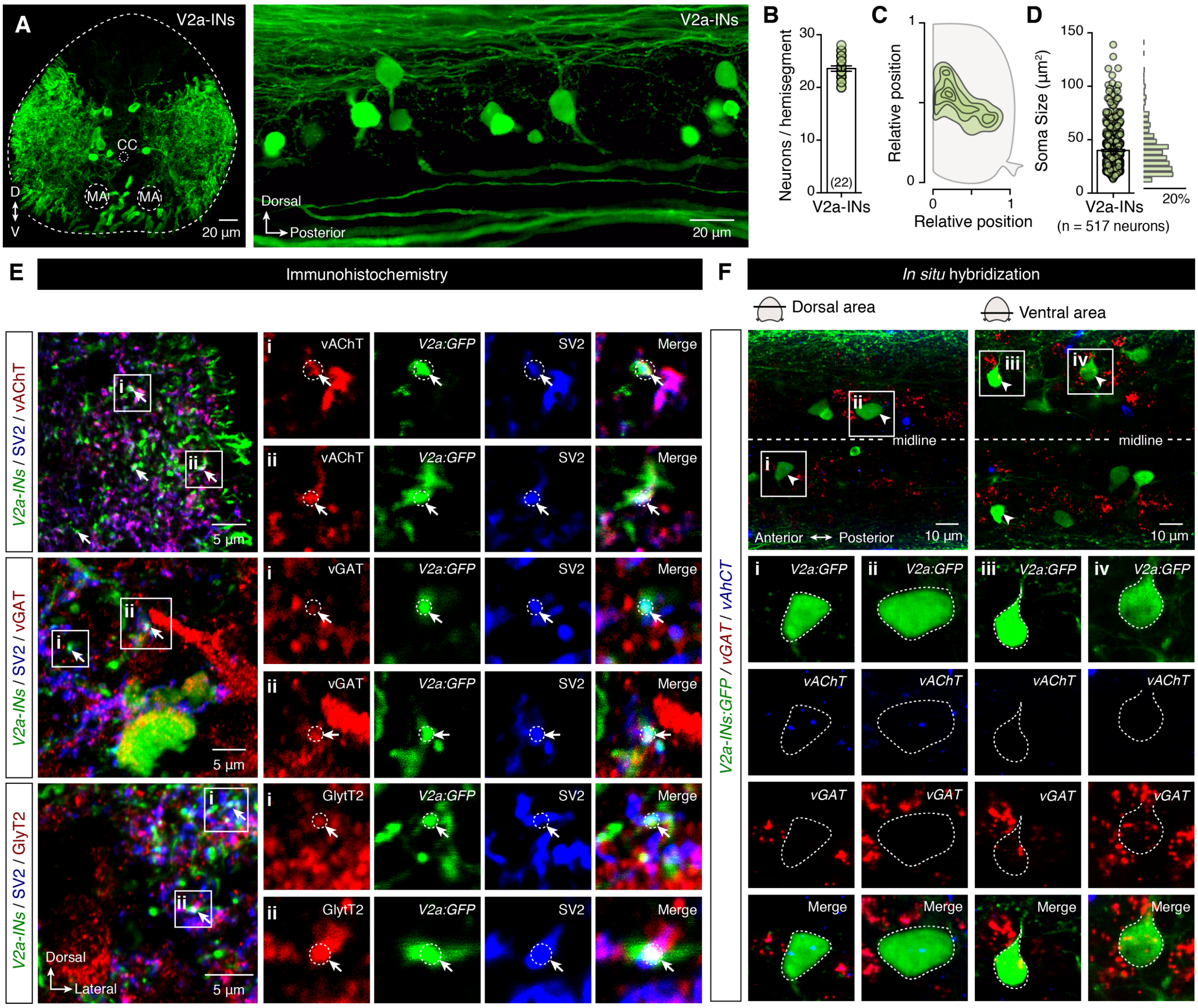

**Figure S4. V2a interneuron analysis**

(A) Transverse section and whole mount adult zebrafish spinal cord showing the distribution of the V2a interneuron population.

(B) Quantification of the number of V2a interneurons in adult spinal cord hemisegment (segment 15).

(C) Spatial distribution of the V2a interneurons with the medio-lateral and dorso-ventral density plot.

(D) Quantification and distribution analysis of the V2a interneuron soma sizes (*n* = 517 neurons).

(E) Confocal photomicrographs of transversal sections of the adult zebrafish spinal cord showing the colocalization of the presynaptic V2a interneuron terminals ($GFP^+/SV2^+$) with the vesicular acetylcholine transporter (vAChT), the vesicular GABA and glycine transporter (vGAT) or with the glycinergic transporter (GlyT2). Arrows indicate the triple co-localization.

(F) Whole mount *in situ* hybridization reveals the co-existence of the different vesicular neurotransmitter transporter mRNAs (*vAChT* or *vGAT*) in the V2a ($GFP^+$) interneurons in the expected relative locations (dorsal, ventral) within the adult zebrafish spinal cord. Arrowheads indicate the double labeled neurons.

Data are presented as mean ± SEM. CC, central canal; MA, Mauthner axon.

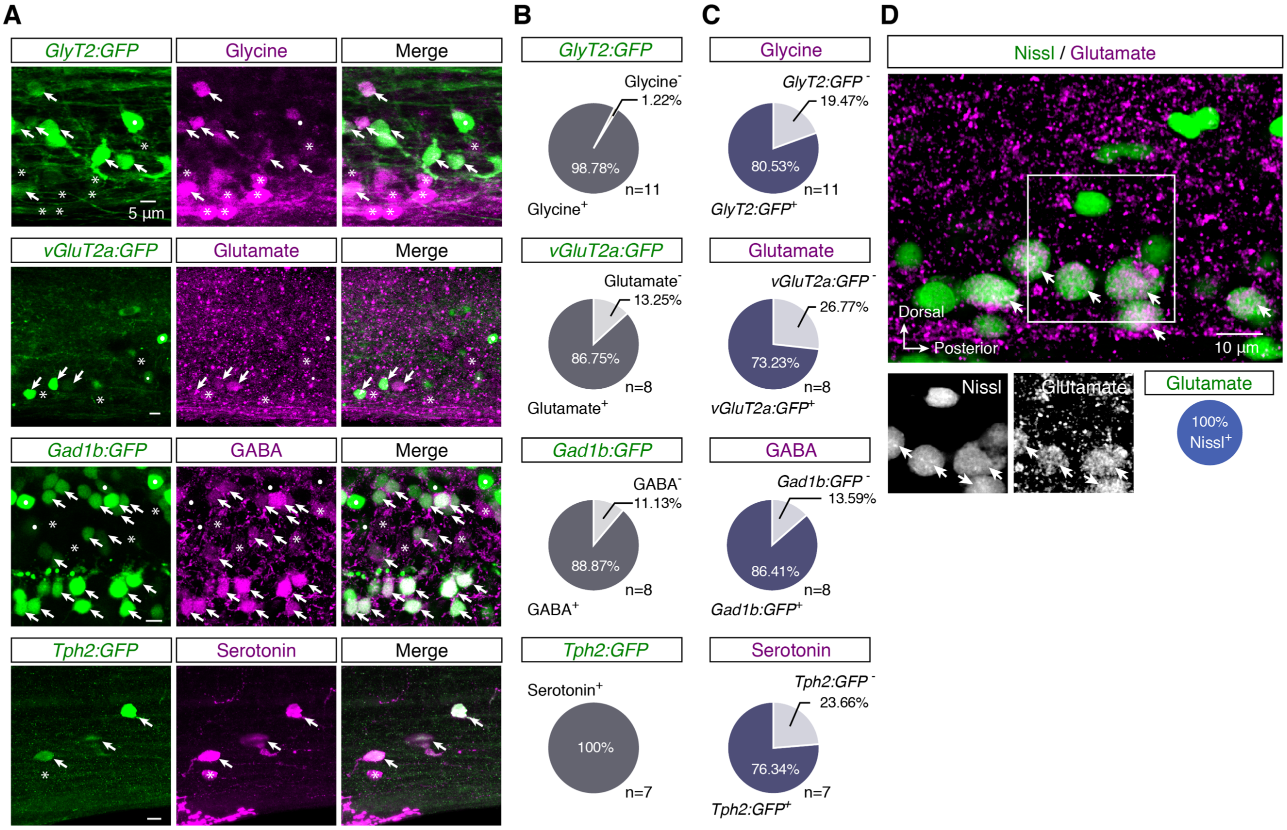


**Figure S5. Transgenic lines capture part of the labeled neuronal populations**

(A) Representative immunofluorescent whole-mount images showing immunostained neurons in the spinal cord of transgenic zebrafish with glycinergic (*GlyT2*), glutamatergic (*vGluT2a*), GABAergic (*Gad1b*) or serotonergic (*Tph2*) neurons genetically labelled (green). Asterisks denote the immunopositive neurons that do not express GFP, dots indicate the GFP expressing neurons that are not immunoreactive and arrows identify double labelled neurons.

(B) Quantification of the percentage of immunolabeled neurons that express GFP.

(C) Quantification of the percentage of GFP$^{+}$ neurons that are labeled with antibodies.

(D) Representative whole mount image of adult zebrafish spinal cord, immunolabeled for glutamate (magenta) with neurons identified by Nissl staining (green). Arrows indicate the double labeled cells.

**TABLE S1**. Antibodies Used[1]

| Antigen | Host | Source | Code | Dilution |
|---|---|---|---|---|
| ***Primary*** | | | | |
| ChAT | Goat | Millipore | AB144P; RRID: AB_2079751 | 1:150 |
| DβH | Rabbit | Millipore | AB1538; RRID: AB_90751 | 1:250 |
| Elav3+4 (HuC/D) | Rabbit | GeneTex | GTX128365; RRID: AB_ | 1:500 |
| GABA | Rabbit | Sigma | A2052; RRID: AB_477652 | 1:2000 |
| GABA | Mouse | From Prof. P. Streit; RRID: AB_2314450 | | 1:700 |
| Glutamate | Rabbit | Sigma | G6642; RRID: AB_259946 | 1:6000 |
| Glycine | Rat | ImmunoSolutions | IG1002; RRID: AB_10013222 | 1:1000 |
| Serotonin | Rabbit | Sigma | S5545; RRID: AB_477522 | 1:4000 |
| Serotonin | Guinea Pig | From Prof. A. Verhofstadt | | 1:2500 |
| TH | Mouse | Millipore | MAB318; RRID: AB_2201528 | 1:800 |
| GFP | Chicken | Abcam | AB13970; RRID: AB_300798 | 1:500 |
| vGAT | Rabbit | Synaptic Systems | 131 002; RRID: AB_887871 | 1:300 |
| GlyT2 | Rabbit | Alomone labs | AGT-012; RRID: AB_11121049 | 1:200 |
| vAChT | Guinea Pig | Millipore | AB1588; RRID: AB_11214110 | 1:1000 |
| SV2 | Mouse | DSHB | SV2; RRID: AB_2315387 | 1:200 |
| ***Secondary*** | | | | |
| Chicken IgY-488 | Goat | ThermoFisher | A-11039; RRID: AB_2534096 | 1:500 |
| Chicken IgY-FITC | Rabbit | ThermoFisher | SA1-9511; RRID: AB_1075130 | 1:500 |
| Goat IgG-568 | Donkey | ThermoFisher | A-11057; RRID: AB_2534104 | 1:500 |
| Goat IgG-488 | Donkey | ThermoFisher | A-11055; RRID: AB_2534102 | 1:500 |
| Goat IgG-647 | Donkey | ThermoFisher | A-21447; RRID: AB_2535864 | 1:500 |
| Guinea Pig IgG-568 | Goat | ThermoFisher | A-11075; RRID: AB_2534119 | 1:500 |
| Mouse IgG-647 | Donkey | ThermoFisher | A-31571; RRID: AB_162542 | 1:500 |
| Mouse IgG-568 | Goat | ThermoFisher | A-11004; RRID: AB_2534072 | 1:500 |
| Mouse IgG-488 | Donkey | ThermoFisher | A-21202; RRID: AB_141607 | 1:500 |
| Rabbit IgG-488 | Donkey | ThermoFisher | A-21206; RRID: AB_2535792 | 1:500 |
| Rabbit IgG-647 | Donkey | ThermoFisher | A-31573; RRID: AB_2536183 | 1:500 |
| Rabbit IgG-568 | Donkey | ThermoFisher | A-10042; RRID: AB_2534017 | 1:500 |
| Rat IgG-550 | Donkey | ThermoFisher | SA5-10027; RRID: AB_2556607 | 1:500 |
| Rat IgG-568 | Goat | ThermoFisher | A-11077; RRID: AB_2534121 | 1:500 |

[1]ChAT, choline-acetyltransferase; DβH, dopamine beta-hydroxylase; GABA, γ-aminobutyric acid; GFP, green fluorescent protein; GlyT2, glycine transporter 2; TH, tyrosine hydroxylase; SV2, synaptic vesicle glycoprotein 2A; vAChT, vesicular acetylcholine transporter; vGAT, vesicular GABA transporter.
